# GATA3 is essential for separating patterning domains during facial morphogenesis

**DOI:** 10.1101/2021.02.17.431667

**Authors:** Makoto Abe, Anthony B. Firulli, Stanley M. Kanai, Kim-Chew Lim, J Douglas Engel, David E. Clouthier

## Abstract

Neural crest cells (NCCs) within the mandibular and maxillary portions of the first pharyngeal arch are initially competent to respond to signals from either region. However, mechanisms that are only partially understood establish developmental tissue boundaries to ensure spatially correct patterning. In the Hinge and Caps model of facial development, signals from both ventral prominences, referred to as the caps, pattern the adjacent tissues while the intervening region, known as the hinge, maintains separation of the mandibular and maxillary domains. One cap signal is GATA3, a member of the GATA family of zinc-finger transcription factors with a distinct expression pattern in the ventral-most part of the mandibular and maxillary portions of the first arch. Here we show that disruption of *Gata3* in mouse embryos leads to hemifacial microsomia, facial bone hypoplasia and syngnathia (bony fusion of the upper and lower jaws). These changes are preceded by gene expression changes in post-migratory NCCs around the maxillomandibular junction (the hinge). GATA3 is thus a crucial component in establishing the network of factors that functionally separate the upper and lower jaws during development.

**Summary Statement:** Loss of *Gata3* leads to BMP-mediated disruption of *Fgf8* expression at the maxillomandibular junction during development, resulting in later fusion of the upper and lower jaws.

## Introduction

Gnathostomes (hinged jaw vertebrates) account for greater than 99% of all vertebrates. Separation of the upper and lower jaws requires establishment and maintenance of gene expression boundaries in the first pharyngeal arch during early embryo development, ensuring that the mandible and maxilla articulate with the skull in a manner that allows the formation of a hinged jaw (Clouthier et al., 2010; Medeiros and Crump, 2012; Neben and Merrill, 2015). Disruption of this process leads to fusion of the upper and lower jaws, a condition called syngnathia (MIM 119550).

One current theory that explains how differential patterning of the first pharyngeal arch is achieved is referred to as the Hinge and Caps model (Depew et al., 2002; Depew et al., 2005; Depew and Compagnucci, 2008). In this model, ectodermal signaling centers (caps) in the ventral (medial) ends of the mandibular and maxillary prominences of the first arch propagate instructive signals dorsally towards the maxillomandibular junction (the hinge), instructing development of the prominences. The maxillomandibular junction separates these two cap domains. As NCCs within the maxillary and mandibular portions of the first arch are at least initially competent to respond to signals normally found in the other half of the arch (Ferguson et al., 2000; Sato et al., 2008; Tavares and Clouthier, 2015), the patterning cues in the hinge domain are equally important to ensure that cap signals do not cross the intervening hinge domain.

Multiple signals establish patterning domains within the mandibular arch of mice and zebrafish, with these domains effectively defining the dorsoventral axis of the arch. These include Endothelin-1 (EDN1) and Bone morphogenetic proteins (BMPs) in the ventral cap, EDN1 in the intermediate domain and Jag/Notch in the dorsal domain (Clouthier et al., 2010; Medeiros and Crump, 2012; Clouthier et al., 2013). Cap signals in the maxillary prominence are also crucial for first arch development and include SHH and ALX (Uz et al., 2010; Dee et al., 2013). However, from a functional standpoint, separation of cap signals by the hinge region (defined as the maxillomandibular junction) is not as well understood. One of the key regulators thus far identified in this process in the mouse is FOXC1, which genetically interacts with *Fgf8* in the hinge region to ensure normal separation of the upper and lower jaw and establishment of the temporomandibular joint (TMJ) (Inman et al., 2013). While loss of *Foxc1* leads to syngnathia, distal cap signals in the mandibular arch are unaffected, indicating that the FOXC1 gene regulatory network acts specifically in the maxillomandibular junction. Other genes acting in the hinge to prevent intrusion of caps signals are not as well established.

GATA3 belongs to the evolutionarily conserved GATA family of zinc finger transcription factors. Mutations in GATA3 are associated with autosomal dominant hypoparathyroidism, sensorineural deafness and renal anomaly (HDR) syndrome (reviewed in (Van Esch and Devriendt, 2001). Targeted deletion of *Gata3* in mice have supported these findings, with deletion resulting in a number of defects, including those that affect cardiac, renal, ocular, craniofacial, sympathetic neuron and immune system development (Pandolfi et al., 1995; Lim et al., 2000; Ho and Pai, 2007; Grote et al., 2008; Hendershot et al., 2008; Maeda et al., 2009; Raid et al., 2009). While the basis of renal, ocular and cardiovascular defects has been characterized, the basis for the craniofacial defects remains unclear.

Here, we demonstrate that loss of *Gata3* disrupts patterning of post-migratory NCCs within pharyngeal arch one due to early disruption of a *Fgf8* GRN in maxillomandibular region. This leads to an intrusion of arch cap signals into the junction, resulting in syngnathia of the upper and lower jaws. Interestingly, the severity of skeletal defects is not bilaterally equal, suggesting that *Gata3* mutant embryos may be a useful model to investigate how facial symmetry is disrupted in human conditions in which facial asymmetry is lost, including hemifacial microsomia (HFM; MIM 164210).

## Experimental Procedures

### Mouse gene and protein nomenclature

All nomenclature for mouse gene and protein names follows official guidelines found at: http://www.informatics.jax.org/mgihome/nomen/gene.shtml

Mouse lines and genotyping. Generation and genotyping of the *Gata3^+/z^* line have been previously described (Lakshmanan et al., 1999). Mice were maintained on a FVB background (Taconic). Genotyping of embryos was conducted using yolk sac DNA and PCR using the primers 5′– TCCTGCGAGCCTGGCTGTCGGA–3′ and 5′–GTTGCCTTGACCATCGATGTT–3′ to detect the wild type allele and 5’-GACACCAGACCAACTGGTA-3’ and 5’-GCATCGAGCTGGGTAATAAC-3’ to detect the *lacZ* allele. Generation of *Gata3^z/z^* embryos for timed matings was conducted by breeding *Gata3^+/z^* animals, with the day when the vaginal plug was observed counted as embryonic day (E) 0.5. All studies were approved by Institutional Animal Care and Use Committee at the University of Colorado Anschutz Medical Campus.

### Skeletal analysis

Analysis of both bone and cartilage development in embryonic (E) 18.5 embryos using Alizarin red and Alcian blue (Ruest et al., 2004) and cartilage analysis in E14.5 embryos using Alcian blue (Clouthier et al., 1998) was performed as previously described. Alizarin red and Alcian blue were obtained from Sigma Aldrich.

### Histology

For histological analysis using hematoxylin and eosin (Sigma Aldrich), embryos were collected and fixed in 4% paraformaldehyde (Fisher Scientific) overnight before being processed through graded ethanols and xylene and then embedded in paraffin as previously described (Tavares et al., 2017). 7 micron sections were cut on a Leica microtome, with staining and subsequent analysis performed as previously described ((Tavares et al., 2017).

### Nerve staining

Analysis of nerve development in E10.5 embryos was performed using a monoclonal anti-neurofilament 160 (NF160) antibody (#N5264, Sigma Aldrich, USA) as previously described (Clouthier et al., 1998). The antibody was used at a dilution of 1:100 and diaminobenzidine (D7304-1SET, Sigma Aldrich, USA) was used as the substrate.

### Sectional cell death and proliferation analysis

Detection of cell proliferation was performed essentially as previously described (Barron et al., 2011). Briefly, pregnant mice were injected intraperitoneally with 200 mg/kg body weight of 5-Ethynyl-2’-deoxyuridine (contained in the Click iT EdU Imaging Kit containing Alexa Fluor 594 as the Alexa Fluor azide (Kit #C10338; Thermo Fisher) 1 h before embryo collection at E9.5. Embryos were collected and fixed in 4% paraformaldehyde (Thermo Fisher Scientific) on ice for 1 hour before rinsing in PBS, dehydrating, and embedding in paraffin. Somite counts were performed before embedding to ensure accurate embryo staging, with three control and three mutant embryos used for the analysis. Embryos were sectioned at 7 microns.

TUNEL analysis was performed using the In Situ Cell Death Detection Kit (Kit #11684795910, Sigma Aldrich) following manufacturer’s recommendations and fluorescein-dUTP. After rinsing slides in PBS, incorporated EdU was detected using the Click iT EdU Imaging Kit according to the manufacturer’s recommendation. 3-4 sections separated by 35-40 microns through the mandibular arch of each wild type or mutant embryo were used for the assay. After staining, sections were counter-stained using DAPI (Sigma) to mark all cell nuclei. Labeled EdU, TUNEL and DAPI cells were then counted, with the TUNEL-positive cells/total cells and EdU-positive cell/total cells calculated. Statistical analysis and figure generation were performed in Prism 9 (GraphPad), with the final graph processed through Adobe Illustrator.

### Whole mount cell death analysis

Whole mount analysis of cell death using TdT-mediated dUTP nick end labeling (TUNEL) was performed as previously described (Abe et al., 2007).

### Whole mount *in situ* hybridization

Gene expression in whole-mount was performed as previously described (Clouthier et al., 1998) using digoxigenin-labeled RNA riboprobes against *Tfap2a* (Williams), *Barx1* (Tissier-Seta et al., 1995), *Bmp4* (Furuta and Hogan 1998), *Dlx2 and Dlx3* (Robinson and Mahon 1994), *Dlx6* (Charite 2001), *Fgf8* (Trumpp et al., 1999), *Hand1* and *Hand2* (Srivastava 1997), *Lhx8* (Tucker et al., 1999), *Msx1* (Thomas 1998), *Pitx1* and *Pitx2* (Liu et al., 2003a).

### *β*-galactosidase staining

To examine *β*-galactosidase (*β*-gal) staining in whole embryos, E8.5, E9.5 and E10.5 *Gata3^z/z^ and Gata3^+/z^* embryos were collected and fixed for 1 hour in 4% paraformaldehyde. Embryo staining and photography was performed as previously described (Ruest et al., 2003). E10.5 embryos were then processed through graded alcohols and embedded transversely in paraffin. 7-micron sections were cut, mounted on plus coated slides, stained with nuclear fast red, coverslipped and photographed (Ruest et al., 2003).

## Results

### Craniofacial defects in *Gata3^z/z^* embryos

To assess the role of GATA3 in craniofacial development, we examined mice in which a *lacZ* reporter cassette was inserted into the *Gata3* gene, resulting in a null allele (Lakshmanan et al., 1999). Since loss of *Gata3* leads to embryonic lethality around embryonic day (E) 13.5 due to noradrenaline deficiency resulting from defects in sympathetic neuron development (Lim et al., 2000), we employed a pharmacological rescue strategy in which pregnant dams were given water containing *α*- and *β*-adrenergic receptor agonists isoproterenol and phenylephrine (Hendershot et al., 2008). Using this approach, *Gata3^z/z^* embryos survived to E18.5, allowing a more complete analysis of bone and cartilage defects. Compared with E18.5 *Gata3^+/+^* embryos (Fig. 1A,C), E18.5 *Gata3^z/z^* embryos presented with retrognathia and had surface ectoderm that almost completely covered the oral opening (Fig. 1B,D). Following skeletal staining, defects in underlying bone and cartilage structures were apparent, including absence of the majority of the proximal mandible and maxilla, with the distal mandible fused with the maxilla (syngnathia) (Fig. 1E,F). The observed bone defects in the upper and lower jaws were not symmetric, with one side always more severe than the other (Fig. 1F,H). This asymmetry resembles hemifacial microsomia (HFM; MIM 164210). Cleft palate was also present (see below).

**Figure 1.**
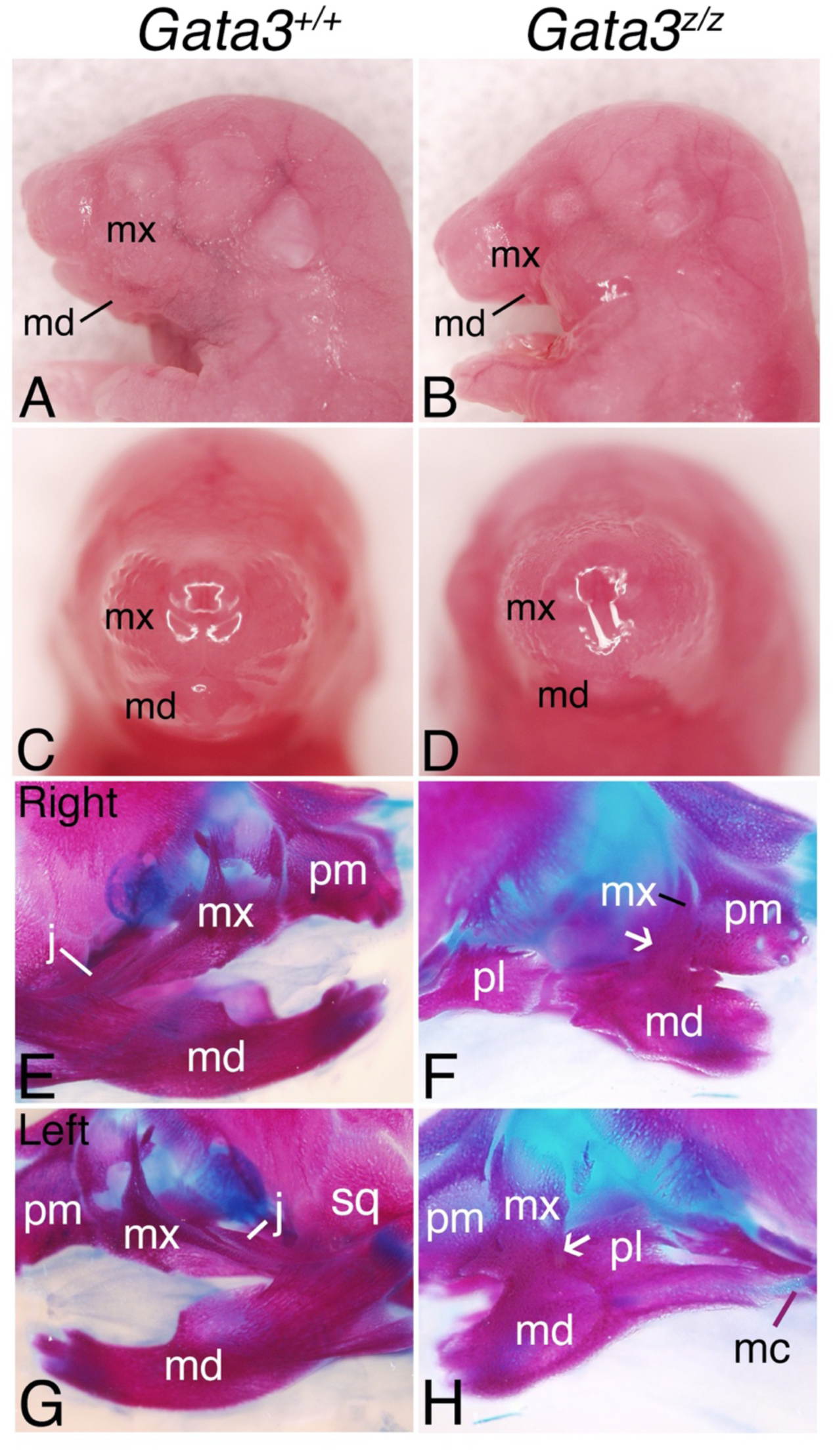
Craniofacial defects in E18.5 *Gata3^z/z^* embryos. **A-D.** Gross views of *Gata3^+/+^* (A, C) and *Gata3^z/z^* (B, D) embryos in lateral (A, B) and ventral (C, D) orientation (n=5 for each group). *Gata3^z^*^/z^ embryos exhibit micrognathia (B, D). **E-H.** Lateral views of skulls from *Gata3^+/+^* (E, G) and *Gata3^z/z^* (F, H) embryos after staining with Alizarin Red (bone) and Alcian Blue (cartilage)(n=5; same embryos as shown in A-D). In *Gata3^z/z^* embryos, the mandible (md) and maxilla (mx) are hypoplastic (most of the proximal bone is missing from both) and fused with each other (arrow in F, H). In addition, the jugal (j) and squamosal (sq) bones are absent in *Gata3^z/z^* embryos and there is an asymmetric loss of Meckel’s’ cartilage (mc; F, H). pl, palatine bones; pm, premaxilla bone.

To define the basis of the syngnathia, we examined earlier skeletal changes in E16.5 *Gata3^+/+^* and *Gata3^z/z^* embryos. Compared with *Gata3^+/+^* embryos (Fig. 2A), the maxilla in *Gata3^z/z^* embryos were present but hypoplastic (Fig. 2B). However, more notable changes were present in the zygomatic arch. The zygomatic arch is composed of the zygomatic process of the maxilla, which articulates with jugal bone, which in turn articulates with the zygomatic process the squamosal bone (Fig. 2A). In *Gata3^z/z^* embryos, the jugal bone failed to extend posteriorly, instead extending caudally and fusing with the mandible (black arrow in Fig. 2B). Mandibular hypoplasia was already evident, including absence of bone in the proximal mandible. Asymmetry in jaw development was also already present, with frontal views of *Gata3^+/+^* (inset in Fig. 2A) and *Gata3^z/z^* (inset in Fig. 2B) embryos clearly showing that the left side of the embryo was more disorganized than the right in *Gata3^z/z^* embryos (n=4).

**Figure 2.**
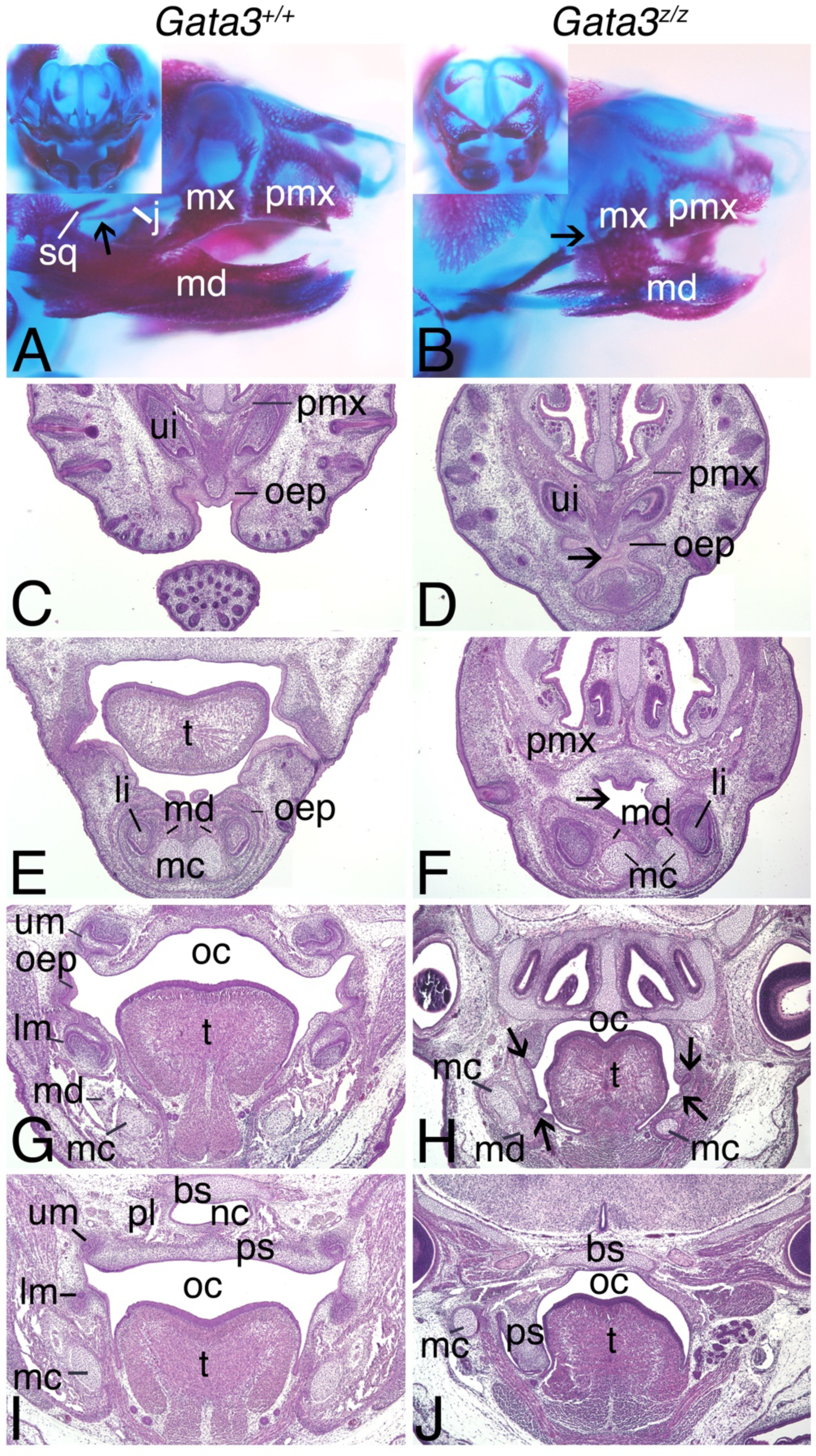
Craniofacial defects in E16.5 *Gata3^z/z^* embryos. **A, B.** Lateral views of *Gata3^+/+^* (A) and *Gata3^z/z^* (B) embryos stained with Alizarin Red and Alcian Blue (n=4 for each genotype). Compared to the mandible in *Gata3^+/+^* embryos (A), the mandible of *Gata3^z/z^* embryos is hypoplastic (B). Also, overall mandibular bone development is more substantial on the right side than the left of *Gata3^z/z^* embryos (compare insets in A and B). In addition, the jugal (j) bone in *Gata3^+/+^* embryos extends anteriorly from the zygomatic process of the maxilla (mx) posteriorly to the zygomatic process of the squamosal bone (sq) (A). In *Gata3^z/z^* embryos, the jugal bone extends towards and fuses to the hypoplastic mandible (md) (arrow in B). **C-J.** Frontal sections of *Gata3^+/+^* (C, E, G, I) and *Gata3^z/z^* (D, F, H, J) embryos stained with hematoxylin and eosin (n=5 for each group). In *Gata3^z/z^* embryos (D, F, H), microstomia is evident (arrow in D, F). In addition, the mandible and Meckel’s cartilage (mc) are asymmetric (F), the tongue (t) is hypoplastic (microglossia) and the upper (um) and lower (lm) molars are arrested at the early bud stage (arrows in H). More proximal sections illustrate that the presence of only one palatal shelf (ps) which has not elevated (J) (3/5 embryos). bs, basisphenoid; li, lower incisor; mc, Meckel’s cartilage; nc, nasal cavity; oep, oral epithelium; pl, palatine bone; pmx, premaxilla; ui, upper incisor.

We next performed histological analysis using frontal sections through the head of E16.5 *Gata3^+/+^* and *Gata3^z/z^* embryos stained with hematoxylin and eosin (H & E) (n=5 for both groups). Compared with *Gata3^+/+^* embryos (Fig. 2C), the lower jaw of *Gata3^z/z^* embryos was fused to the upper jaws, with oral ectoderm almost completely obstructing the oral opening, resulting in microstomia (Fig. 2D).

More proximally in the oral cavity, the lower jaw of *Gata3^+/+^* embryos appeared symmetric (Fig. 2E), whereas the lower jaw of *Gata3^z/z^* embryos was shifted to the left and again showed microstomia (Fig. 2F). In these mutants, the mandible, Meckel’s cartilage and incisors appeared dysmorphic. In addition, Meckel’s cartilage was not fused at its symphysis (Fig. 2F) and the tongue was hypoplastic, with a disorganization of the connective tissue and muscle fibers (Fig. 2H) compared with the organization observed in the tongues of *Gata3^+/+^* embryos (Fig. 2G). While the upper and lower incisors showed some variation in size in *Gata3^z/z^* embryos, overall development appeared normal (Fig. 1F,H; Fig. 2F). In contrast, both upper and lower molar development appeared arrested at the early bud stage (Fig. 2H, arrows). Cleft palate was also present in all *Gata3^z/z^* embryos; in sectioned embryos, the right palatal shelf was not elevated while the with the left shelf was absent (Fig. 2J; n=3), while other embryos showed two non-elevated palatal shelves (n=2; data not shown). Most proximal jaw structures were absent, including the mandibular condyles and the temporomandibular joint (data not shown). Other changes included hypoplasia of the tympanic ring and gonial bone and absence of the alisphenoid bone and most of the squamosal bone (data not shown; also apparent in Fig. 2F,H). The middle ear ossicles (malleus, incus, and stapes) appeared normal (data not shown).

### Early developmental defects in *Gata3^z/z^* embryos

Disruption in early pharyngeal arch patterning signals often also affect cranial ganglia development. When *Gata3^+/+^* embryos were stained in whole mount with an antibody against NF160, a neuronal marker, the ophthalmic, maxillary and mandibular branches of the trigeminal ganglia and the facial (VII) nerve were normal in appearance and projection (Fig. 3A). In contrast, a majority of the maxillary branch of the trigeminal ganglia in stained *Gata3^z/z^* embryo was absent, though a small piece of nerve tissue remained below the optic placode (Fig. 3B). In addition, the facial nerve (VII) showed numerous aberrant fasciculations. These two defects were observed on both sides (n=5), though individual fasciculations varied between sides and embryos.

**Figure 3.**
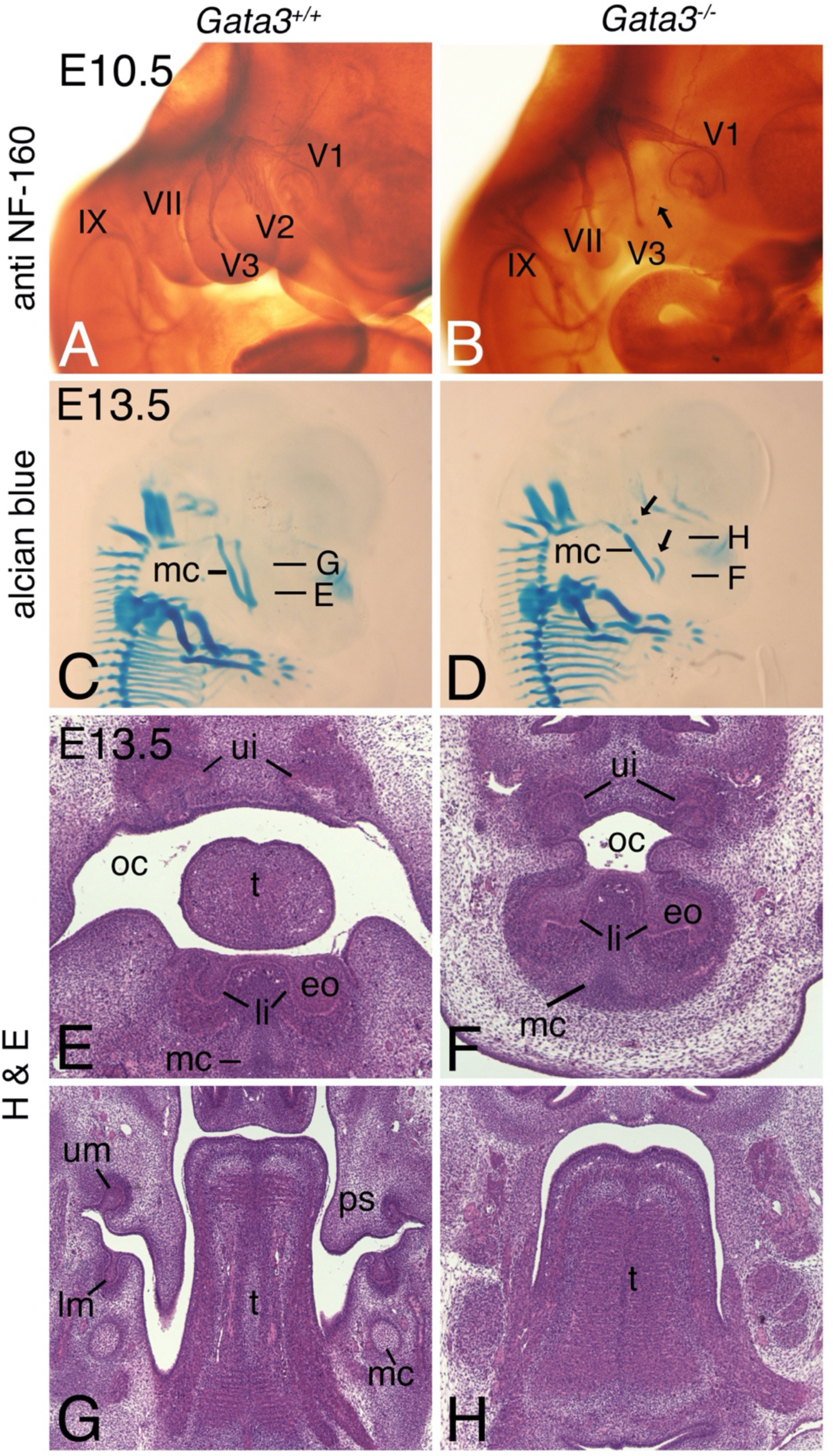
Early developmental defects in *Gata3^z/z^* embryos. **A, B.** Lateral views of E10.5 *Gata3^+/+^* (A) and *Gata3^z/z^* (B) embryos processed for whole mount immunohistochemistry with an antibody against the neurofilament protein 160 (NF-160) (n=5). While the ophthalmic (V1), maxillary (V2) and mandibular (V3) branches of the trigeminal (V) ganglion are observed in *Gata3^+/+^* embryos (A), the V1 is absent in *Gata3^z/z^* embryos, though a small bundle of distal nerve is observed (B, arrow). Numerous aberrant fasciculations are also present along facial nerve (VII). **C, D.** Lateral views of E13.5 *Gata3^+/+^* (C) and *Gata3^z/z^* (D) embryos stained with Alcian Blue (cartilage) show that Meckel’s cartilage (mc) in *Gata3^z/z^* embryos has a gap on each side, though the left gap is more prominent (gap denoted by black arrows) (D) (n=4 for each group). Letters in panels denote the plane of section shown in panels E-H. **E-H.** Frontal sections through the head of E13.5 *Gata3^+/+^* (E, G) and *Gata3^z/z^* (F, H) embryos stained with hematoxylin and eosin (n=4 for each group). While the oral cavity is smaller in *Gata3^z/z^* embryos (F) compared with *Gata3^+/+^* embryos (E), the lower incisors (li) appear similar between the two genotypes. Further proximal in the oral cavity, the tongue (t) is smaller in *Gata3^z/z^* embryos (H) compared to *Gata3^+/+^* embryos (G). In addition, the palatal shelves (ps) and developing upper (um) and lower (lm) molars in *Gata3^z/z^* embryos are absent (H). IX, glossopharyngeal nerve; eo, enamel organ; ui, upper incisors.

By E13.5, Meckel’s cartilage was apparent in *Gata3^+/+^* embryos, extending from the symphysis of the mandible to the forming temporomandibular joint (Fig. 3C). In E13.5 *Gata3^z/z^* embryos, Meckel’s cartilage was present but curved downward at on the distal end (Fig. 3D). In addition, the central portion of Meckel’s cartilage was absent on the left side (area between black arrows, Fig. 3D; n=4). In hematoxylin and eosin (H & E)-stained frontal sections through the head of E13.5 embryos, incisors were present in both *Gata3^+/+^* (Fig. 3E) and *Gata3^z/z^* (Fig. 3F) embryos, though lower incisors were slightly dysmorphic in *Gata3^z/z^* embryos (n=4). Further, the microstomia observed in E16.5 *Gata3^z/z^* embryos (Fig. 2D, F) was already apparent at this earlier age, with only a small portion of the oral cavity separating the upper and lower incisors (Fig. 3F). At the level of the molars, upper and lower molars were present in *Gata3^+/+^* embryos (Fig. 3G), with downward-oriented palatal shelves and Meckel’s cartilage also present. In *Gata3^z/z^* embryos, molar tooth buds were not obvious (Fig. 3H). In addition, palatal shelves were either hypoplastic (n=2; data not shown) or not present (Fig. 3H; n=2).

### Defects in early NCCs in *Gata3^z/z^* embryos

Bone hypoplasia/absence in the mandibular and maxillary prominences of the first pharyngeal arch and the absence of the maxillary branch of the trigeminal nerve in *Gata3^z/z^* embryos are suggestive of defects in NCC specification, migration or differentiation. To assess these possibilities, we examined expression of the NCC marker *Tfap2*, a transcription factor involved in multiple facets of cranial NCC (Brewer et al., 2004). In E8.5 *Gata3^+/+^* embryos, *Tfap2a* expression was observed along the neuroepithelium of the posterior midbrain and hindbrain, with expression also present caudal to the preotic sulcus (pos) (Fig. 4A,B). While *Tfap2a* expression was present in the staining in the posterior midbrain and hindbrain of E8.5 *Gata3^z/z^* embryos, the relative level was reduced, while expression caudal to the pos was more dramatically reduced (Fig. 4C,D).

**Figure 4.**
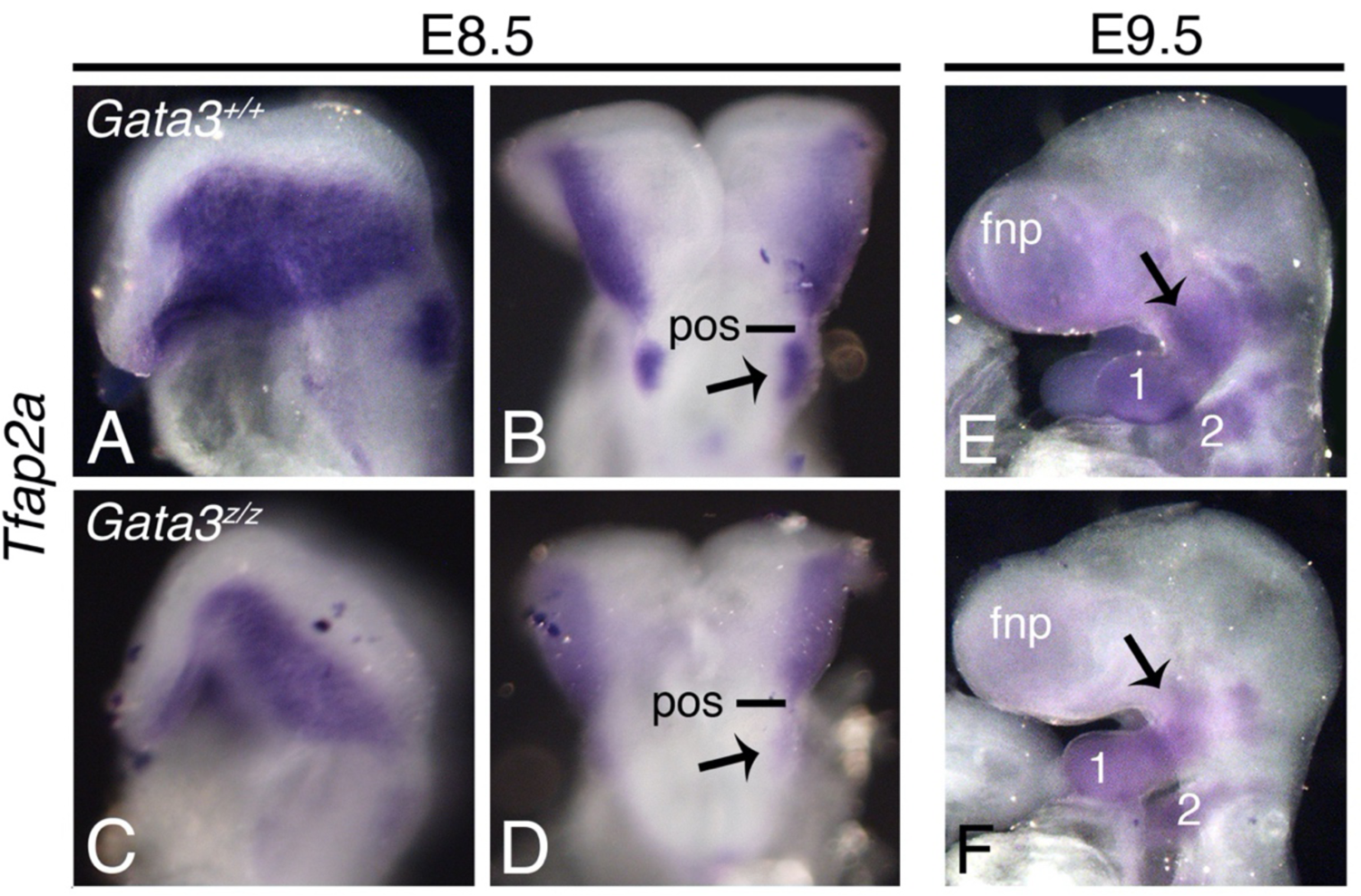
Expression of neural crest marker gene *Tfap2* in *Gata3^z/z^* embryos. Whole mount in situ hybridization analysis in E8.5 (A-D) and E9.5 (E, F) *Gata3^+/+^* (A, B, E) and *Gata3^z/z^* (C, D, F) embryos (n=4 for each group). **A-D.** *Tfap2* expression is similar in E8.5 *Gata3^+/+^* (A, B) and *Gata3^z/z^* (C, D) embryos, though expression in *Gata3^z/z^* embryos is only weakly expressed past the post-otic sulcus (pos; arrow). **E, F.** *Tfap2* expression in E9.5 *Gata3^+/+^* (K) and *Gata3^z/z^* (L) embryos is observed extending in streams from the posterior midbrain/hindbrain to the pharyngeal arches, though the expression appears weaker in *Gata3^z/z^* embryos, including around the maxillomandibular junction (compare arrows in E and F). 1, first pharyngeal arch; 2, second pharyngeal arch; fnp, frontonasal process.

By E9.5, *Tfap2a* expression in *Gata3^+/+^* embryos was broadly observed in the mandibular portion of arch one, extending towards the maxillary prominence (black arrow in Fig. 4E). A similar expression was observed in E9.5 *Gata3^z/z^* embryos, though overall staining appeared reduced, including the area near the maxillary prominence (black arrow, Fig. 4F). These findings indicate that while there was not a major disruption in NCC migration in *Gata3^z/z^* embryos, eighter fewer NCCs are reaching the arches or normal gene expression within the NCCs is disrupted.

### Cell death and proliferation in the mandibular arch of *Gata3^z/z^* embryos

While fewer migrating NCCs could account for pharyngeal arch hypoplasia and loss of bone structures in E18.5 embryos, we also examined whether there was a difference in early proliferation or apoptosis in the NCC-derived mesenchyme within the mandibular arch at E9.5, a time at which NCC migration is complete and arch patterning is beginning. Whole mount terminal deoxynucleotidyl transferase biotin-dUTP nick end labeling (TUNEL) analysis in E9.5 embryos revealed that the relative incidence of cell death between *Gata3^+/+^* (Fig. 5A) and *Gata3^z/z^* (Fig. 5B) embryos appeared similar. TUNEL analysis on sections through the mandibular arch mesenchyme of E9.5 embryos also did not show a statistically significant difference in between *Gata3^+/+^* and *Gata3^z/z^* embryos (Fig. 5C). Similarly, there was not a statistically significant difference in proliferation in the mandibular arch mesenchyme between *Gata3^+/+^* and *Gata3^z/z^* embryos (Fig. 5D). However, overall cell number in the mandibular arch mesenchyme of *Gata3^z/z^* mutants was significantly reduced compared with *Gata3^+/+^* embryos, with one possibility being that fewer NCCs populated the mandibular arch of *Gata3^z/z^* embryos.

**Figure 5.**
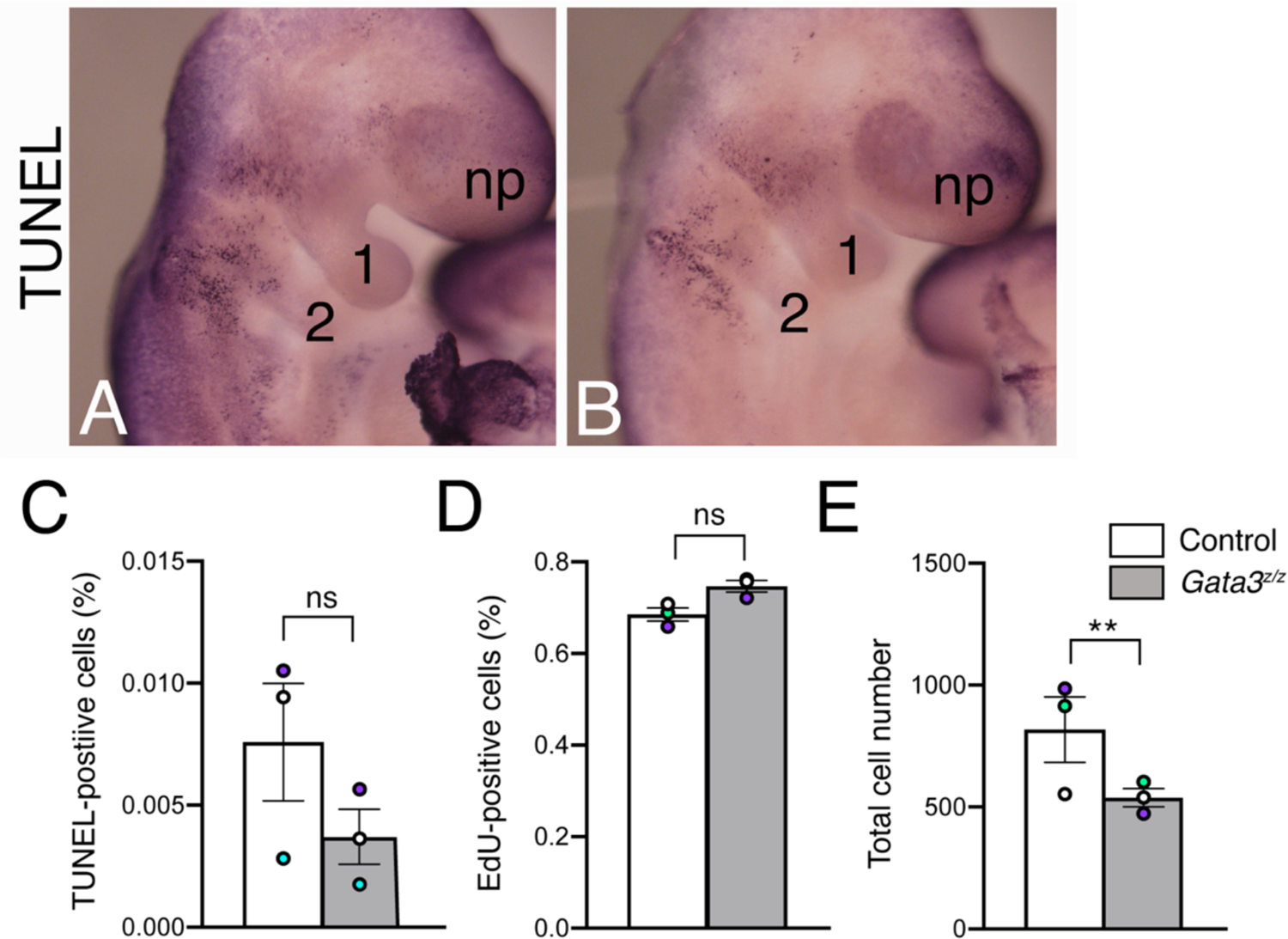
Change in total mesenchyme cell number in the mandibular arch of *Gata3^z/z^* embryos. A, B. Representative lateral views of E9.5 *Gata3^+/+^* (A) and *Gata3^z/z^* (B) embryos processed for whole mount terminal deoxynucleotidyl transferase biotin-dUTP nick end labeling (TUNEL) staining (total examined for each group = 3. The incidence of cell death is qualitatively similar between the two genotypes. **C-E.** Quantification of cell death (C), proliferation (D) and cell number (E) within the mandibular arch mesenchyme in E9.5 *Gata3^+/+^* and *Gata3^z/z^* embryos (n=3). Three sections through the mandibular arch (separated by 30-35 microns) of three *Gata3^+/+^* or *Gata3^z/z^* embryos were processed for analysis. Data is presented as an average +/- s.e.m. per genotype, with each colored circle representing a single embryo. Statistics on the underlying nine values for each group were performed using Prism and an unpaired two-tailed t test. **C.** TUNEL-positive cells following TUNEL analysis of sectioned embryos. Differences between *Gata3^+/+^* and *Gata3^z/z^* embryos were not statistically significant (p=0.18). **D.** Quantification of 5-ethynyl-2’-deoxyuridine (EDU)-positive cells in the mandibular arch on the same sections as those used in C. Differences in the number of proliferative cells were not statistically significant between *Gata3^+/+^* and *Gata3^z/z^* embryos (p=0.12). **E.** The total number of cells in the mandibular arch of *Gata3^z/z^* embryos is significantly less than the total number in *Gata3^+/+^* embryos (p=0.008). n.s. not significant, **, p<0.01.

### GATA3 disrupts gene expression surrounding the hinge region

We have previously shown that *Gata3* is expressed in the ventral domain of the mandibular arch (the cap of the mandibular arch) and the lambdoidal junction between the medial and lateral nasal prominences (the cap of the maxillary prominence) by E9.5, with this pattern more prominent by E10.5 (Ruest et al., 2004). While the mandibular arch expression domain of *Gata3* corresponds to the *Hand2* expression domain, *Hand2* expression is induced by EDNRA signaling (Clouthier et al., 2000; Charité et al., 2001; Ruest et al., 2004), while *Gata3* expression is largely independent (Ruest et al., 2004); in fact, in E10.5 embryos, ventral *Hand2* expression in *Gata3^z^*^/z^ embryos expands into the dorsal mandibular arch along the rostral half of the arch, approaching the maxillomandibular junction (Ruest et al., 2004). These findings suggest that GATA3 may regulate a gene regulatory network (GRN) that maintains gene expression around the maxillomandibular junction, in part by preventing encroachment of cap gene expressing into the junction, thus allowing for development of distinct upper and lower jaws. This could explain, in part, the observed phenotype of *Gata3^z/z^* embryos

To further investigate this possibility, we examined the expression of genes known to pattern the dorsal mandibular arch. We first examined *Pitx2* expression, as PITX2 acts in a dosage-dependent manner to regulate the expression of a number of other genes around the maxillomandibular junction (Liu et al., 2003b). In E9.5 *Gata3^+/+^* embryos, *Pitx2* expression extended along the ectoderm from the maxillary prominence to the rostral half the mandibular arch (Fig. 6A; extent denoted by arrows) (Liu et al., 2003b). A similar expression pattern was observed in E9.5 *Gata3^z/z^* embryos, though both the maxillary and mandibular prominences appeared smaller than those observed in *Gata3^+/+^* embryos (Fig. 6B; extent denoted by arrows). In E10.5 *Gata3^+/+^* and *Gata3^z/z^* embryos, the *Pitx2* expression pattern was similar to that observed at E9.5 (Fig. 6C,D; extent denoted by arrows) (Liu et al., 2003b), though as observed at E9.5, both prominences appeared smaller in *Gata3^z/z^* embryos.

**Figure 6.**
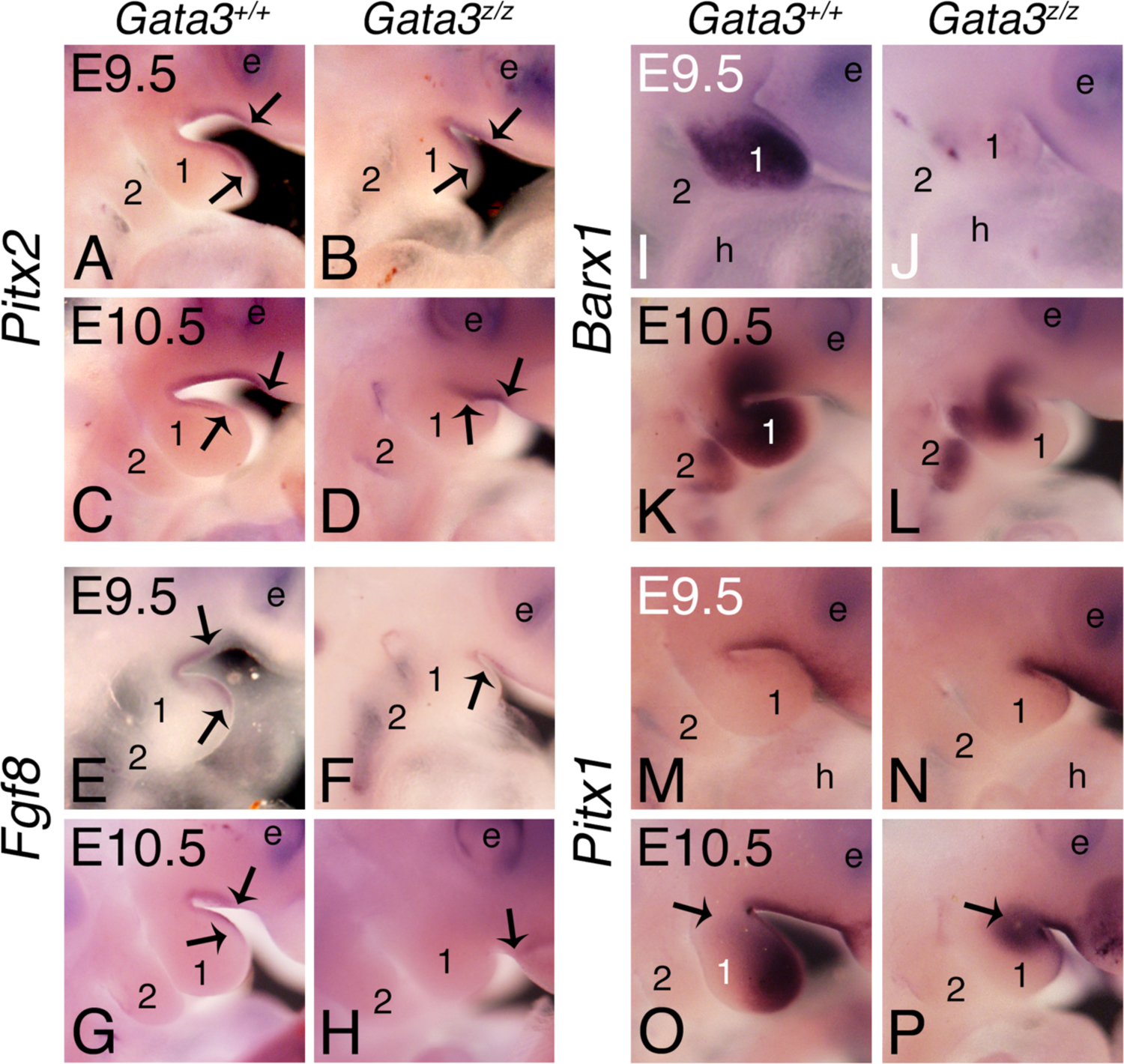
A PITX2-mediated gene regulatory network is disrupted by loss of GATA3. Lateral views of *Gata3^+/+^* (A,C,E,G,I,K,M,O) and *Gata3^z/z^* (B,D,F,H,J,L,N,P) embryos following whole mount in situ hybridization (n=4 for each group). **A-D.** *Pitx2* is expressed along the ectoderm of the maxillary and mandibular (1) prominences and spans the maxillomandibular junction in *Gata3^+/+^* (A, C) and *Gata3^z/z^* (B, D) embryos at both E9.5 (A, B) and E10.5 (C, D)(boundary denoted by arrows), though the mandibular arch is smaller in *Gata3^z/z^* embryos (D; arrows)**. E-H.** *Fgf8* is expressed along the ectoderm spanning the maxillomandibular junction in E9.5 and E10.5 *Gata3^+/+^* embryos (E, G; boundary denoted by arrows). While *Fgf8* expression in *Gata3^z/z^* embryos is weakly expressed around the maxillomandibular junction at E9.5 (F; arrows), expression is only observed near the lambdoidal junction at E10.5 (H, arrow). **I, L.** *Barx1* is expressed in the mandibular arch mesenchyme of E9.5 *Gata3^+/+^* embryos (I) but is almost completely absent from the arch mesenchyme of E9.5 *Gata3^z/z^* embryos (J). At E10.5, *Barx1* is expressed in the mesenchyme of the maxillary and mandibular portions of arch one and along the maxillomandibular junction (K). In E10.5 *Gata3^z/z^* embryos, *Barx1* expression is less pronounced and centered around the maxillomandibular junction (L). **M-P.** *Pitx1* is expressed in the ectoderm of the maxillary prominence and the maxillomandibular junction in E9.5 *Gata3^+/+^* (M) and *Gata3^z/z^* (N) embryos. In E10.5 *Gata3^+/+^* embryos, *Pitx1* expression in the ectoderm expression persists and is now present in the mandibular arch mesenchyme, though expression weakens closer to the maxillomandibular junction (O; arrow). In E10.5 *Gata3^z/z^* embryos (P), *Pitx1* is also expressed along the maxillary ectoderm but mesenchyme expression is confined to the area around the maxillomandibular junction (arrow).

We next examined expression of the PITX2 target gene *Fgf8* (Liu et al., 2003b). At E9.5, *Fgf8* expression in *Gata3^+/+^* embryos was observed along the ectoderm of the maxillary prominence and rostral half of the mandibular arch, spanning the maxillomandibular junction (Fig. 6E; extent denoted by arrows) (Trumpp et al., 1999; Liu et al., 2003b). The *Fgf8* expression pattern was similar in E9.5 *Gata3^z/z^* embryos (Fig. 6F), though was less pronounced and appeared broader on the maxillary prominence than observed in *Gata3^+/+^* embryos. By E10.5, *Fgf8* expression in *Gata3^+/+^* embryos was again observed along the ectoderm of the maxillary and mandibular prominences (Fig. 6G, extent denoted by arrows) (Liu et al., 2003b). In contrast, *Fgf8* expression in E10.5 *Gata3^z/z^* embryos expression was undetectable by ISH in the first arch ectoderm (Fig. 6H). This absence suggests that changes in *Fgf8* expression are unrelated to PITX2 activity.

We next examined *Barx1*, a prominent FGF8 target whose expression is sensitive to reduction in *Fgf8* expression (Trummp et al., 1999). In E9.5 *Gata3^+/+^* embryos, strong *Barx1* expression was observed in the dorsal and intermediate domains of the mandibular arch mesenchyme (Fig. 6I) (Clouthier et al., 2000) but was almost absent in E9.5 *Gata3^z/z^* embryos (Fig. 6J). At E10.5, *Barx1* expression in *Gata3^+/+^* embryos was observed in the mesenchyme of the mandibular and maxillary portions of arch one (spanning the maxillo-mandibular junction) and in the mesenchyme of arch two (Fig. 6K) (Clouthier et al., 2000). In E10.5 *Gata3^z/z^* embryos, expression in arch two was similar to that of *Gata3^+/+^* embryos, though arch one expression was reduced in both the mandibular and maxillary prominences to an area immediately surrounding the maxillomandibular junction (Fig. 6L).

*Pitx1* expression is sensitive to *Pitx2* gene dosage (Liu et al., 2003b), though its expression relies also on *Fgf8* expression (Trumpp et al., 1999). At E9.5, *Pitx1* expression appeared along the ectoderm of the maxillary prominence, where it extended just past the maxillomandibular junction in *Gata3^+/+^* (Fig. 6M) (Lanctôt et al., 1997) and *Gata3^z/z^* embryos (Fig. 6N). By E10.5, *Pitx1* expression in *Gata3^+/+^* embryos remained along the ectoderm of the maxillary and mandibular arches while also expanding into the mandibular arch mesenchyme (Fig. 6O) (Lanctôt et al., 1997; Liu et al., 2003b). In contrast, while ectoderm expression of *Pitx1* was unchanged in E10.5 *Gata3^z/z^* embryos, *Pitx1* expression in the mandibular arch mesenchyme was shifted dorsally, localizing around the maxillomandibular junction (Fig. 6P). Thus, loss of *Gata3* disrupts a FGF8 GRN that includes *Barx1* and *Pitx1*, with expression of these genes either greatly reduced or shifted more dorsally.

The presence of *Fgf8* expression from the maxillomandibular junction establishes the dorsal identity in the first arch, though also relies on the exclusion of BMP signaling (Liu et al., 2003b). In E9.5 *Gata3^+/+^* embryos, prominent *Bmp4* expression was observed on the ectoderm of the dorsal maxillary and mandibular portions of arch one (boundaries denoted black arrows in Fig. 7A) (Liu et al., 2005). Importantly, there appeared little overlap of *Bmp4* (Fig. 7A) and *Fgf8* expression (Fig. 6A). In E9.5 *Gata3^z/z^* embryos, *Bmp4* expression was observed on the ectoderm of the maxillary and mandibular portions of arch one (arrow), including the ectoderm of the maxillomandibular junction (Fig. 7B). In E10.5 *Gata3^+/+^* embryos, ectodermal expression of *Bmp4* remained excluded from the maxillomandibular junction region (Fig. 7C) (Liu et al., 2005), whereas *Bmp4* expression spanned this junction in E10.5 *Gata3^z/z^* embryos (Fig. 7D).

**Figure 7.**
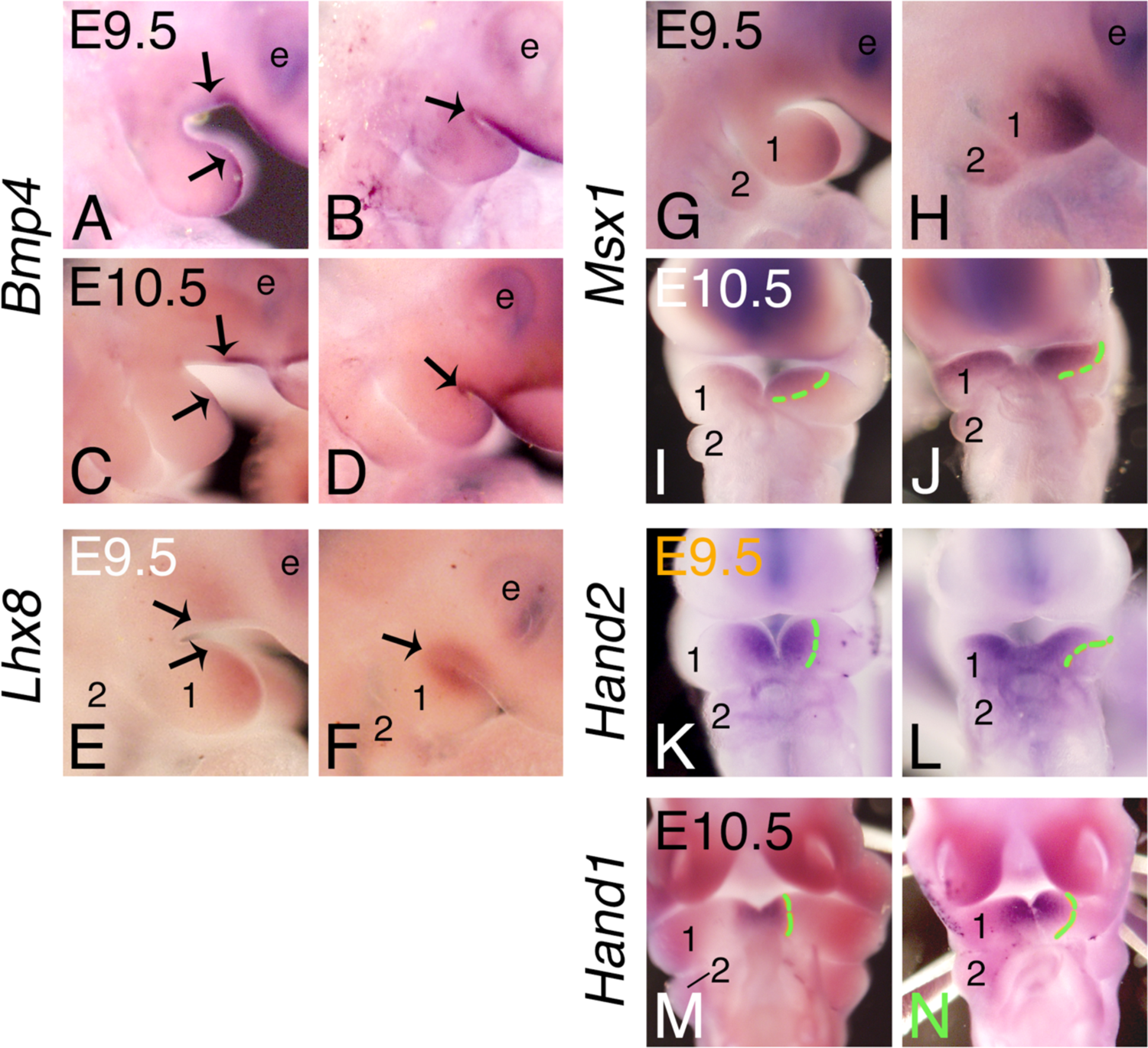
A BMP gene regulatory network is disrupted in *Gata3^z/z^* embryos. **A-J**. Lateral (A-H) and frontal (I-N) views of *Gata3^+/+^* (A,C,E,G,I,K,M) and *Gata3^z/z^* (B,D,F,H,J,L,N) embryos at E9.5 and E10.5 after whole-mount in situ hybridization (n=4 for each group). **A-D.** *Bmp4* is expressed along the ectoderm of the mandibular and maxillary prominences in E9.5 (A) and E10.5 (C) *Gata3^+/+^* embryos but is excluded from the *Fgf8* expression domain along the ectoderm of the maxillomandibular junction (shown in Fig. 6E, G). In *Gata3^z/z^* embryos, *Bmp4* expression spans the maxillomandibular junction at both E9.5 (B) and E10.5 (D). **E, F.** In E9.5 *Gata3^+/+^* embryos *Lhx8* is expressed in the mesenchyme of the maxillary and mandibular prominences of the first arch but is excluded from the maxillomandibular junction (borders denoted by arrows). (E). In E9.5 *Gata3^z/z^* embryos, *Lhx8* expression is confined to the mesenchyme surrounding the maxillomandibular junction (arrow)(F). **G-J.** *Msx1* is expressed in the ventral mandibular arch mesenchyme of E9.5 (G) and E10.5 (I) *Gata3^+/+^* embryos (denoted at E10.5 by a green dashed line in I) and weakly in the maxillary prominence. In E9.5 (I) and E10.5 (J) *Gata3^z/z^* embryos *Msx1* is expressed in the ventral mandibular arch mesenchyme, though the expression boundary extends dorsally toward the maxillomandibular junction (denoted by a green dashed line in J). Weak expression in still present in the maxillary prominence. **K, L.** *Hand2* expression is restricted to the ventral domain in E9.5 *Gata3^+/+^* embryos (K; denoted by the green dashed line). In E9.5 *Gata3^z/z^* embryos, *Hand2* is expressed in the distal domain, though expression extends on the dorsorostrally (L; denoted by the green dashed line). **M, N.** *Hand1* expression is confined to the ventral mandibular arch in E10.5 *Gata3^+/+^* (M) and *Gata3^z/z^* (N) embryos.

We next examined expression of two BMP4-induced genes whose expression is upregulated when *Bmp4* expression is upregulated (Liu et al., 2003b; Bonilla-Claudio et al., 2012). In E9.5 *Gata3^+/+^* embryos, *Lhx8* expression was observed in the ventral mesenchyme of the mandibular arch and in the mesenchyme of the of the maxillary prominences but not in the mesenchyme near the maxillomandibular junction (Fig. 7E; boundary of expression denoted by arrows) (Tucker et al., 1999). In E9.5 *Gata3^z/z^* embryos, *Lhx8* expression shifted to the area around the maxillomandibular junction (Fig. 7F) in a pattern similar to aberrant *Bmp4* expression in *Gata3^z/z^* embryos (Fig. 7B). A similar shift in gene expression was observed for *Msx1*, which in E9.5 *Gata3^+/+^* embryos was located in the ventral mandibular arch mesenchyme (Fig. 7G, I) (Tucker et al., 1999). In E9.5 *Gata3^z/z^* embryos, *Msx1* mesenchyme expression was observed in the mesenchyme adjacent the ectoderm along the maxillary and mandibular prominences, including around the maxillomandibular junction (Fig. 7H, J). In the mandibular arch, this expansion was confined to the rostral arch in a pattern that again represented an area adjacent to increased ectodermal *Bmp4* expression (Fig. 7B). *Hand2* expression in E9.5 *Gata3^+/+^* embryos was confined ventrally (Fig. 7K) (Clouthier et al., 2000) while expression in E9.5 *Gata3^z/z^* embryos extended dorsally along the rostral half of the arch (Fig. 7L). This shift in expression was similar to the *Msx1* expression pattern (Fig. 7J) and one we have previously observed for *Hand2* expression in E10.5 *Gata3^z/z^* embryos (Ruest et al., 2004).

These changes in the BMP GRN could reflect a gain in BMP signaling in the maxillomandibular junction area and/or inappropriate patterning of ventral mandibular arch cells due closer apposition of ventral cells with the maxillomandibular junction. The latter would occur if there was an absence or reduction in dorsal domain cells. To examine this question, we determined whether the expression of *Hand1*, one of the ventral-most markers in the mandibular arch, was altered. In E10.5 *Gata3^+/+^* and *Gata3^z/z^* embryos, *Hand1* expression was confined to the ventral arch mesenchyme (Fig. 7M, N). This suggests that the expansion of ventral and intermediate gene expression into maxillomandibular junction is not simply due to proximity of the ventral cap to the maxillomandibular junction.

### Loss of GATA3 results in expansion of the *Gata3* expression domain

The expansion of ventral gene expression towards the maxillomandibular junction illustrates that the absence of GATA3 results in a dorsal expansion of a ventral GRN. Since *Gata3* is a ventral (cap) marker, we examined whether targeted inactivation of *Gata3* also affected the transcriptional control of *Gata3* itself. To accomplish this, we took advantage of the *lacZ* gene knocked-in to the *Gata3* locus. At E8.5, *β*-galactosidase (*β*-gal) activity was observed in pharyngeal arches of both *Gata3^+/z^* (Fig. 8A) and *Gata3^z/z^* (Fig. 8B) embryos. Additional staining in *Gata3^z/z^* embryos was observed extending from the neural tube to the arches/circumpharyngeal region (white arrows in Fig. 8A,B). At E9.5, *β*-gal staining within the arches was observed in the ventral domain of the pharyngeal arches.

**Figure 8.**
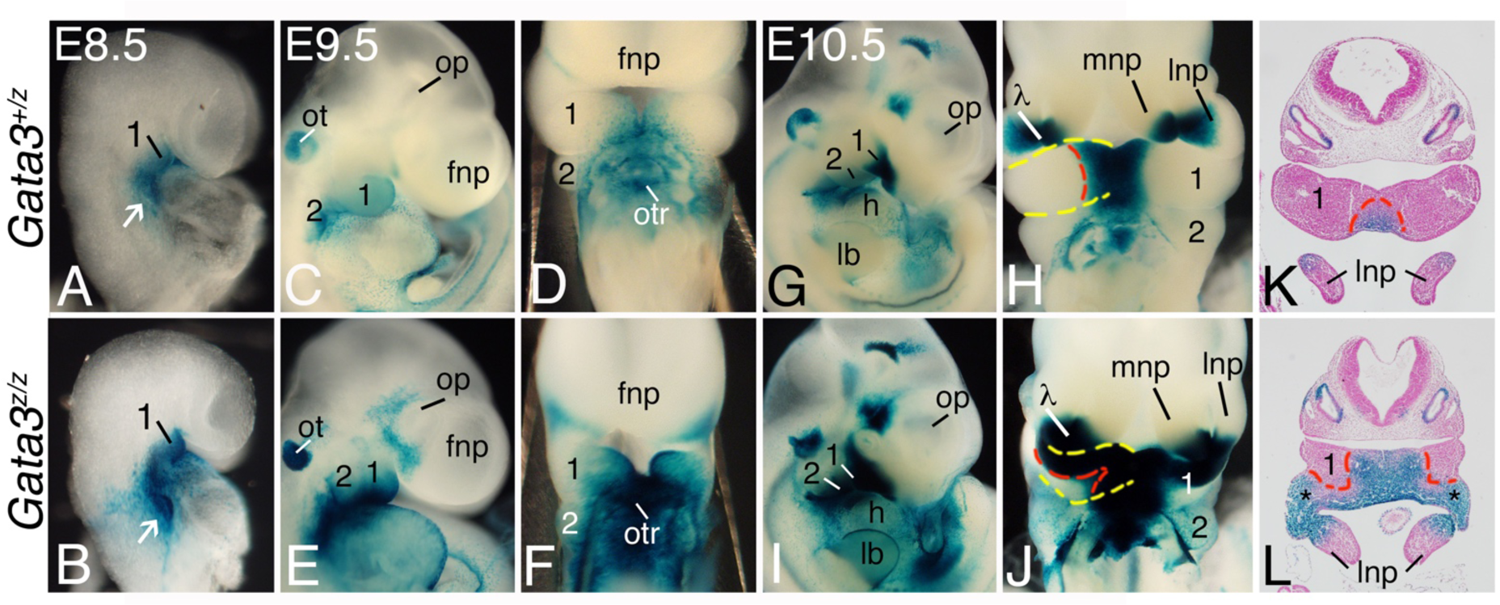
*Gata3* transcription expands in *Gata3^z/z^* embryos. β-galactosidase (β-gal) activity in *Gata3^+/z^* (A,C,D,G,H,K) and *Gata3^z/z^* (B,D,F, H,J,L) embryos at E8.5, E9.5, and E10.5 (n=6 at each time point). **A, B.** Lateral views of E8.5 *Gata3^+/z^* (A) and *Gata3^z/z^* (B) embryos showing β-gal activity in the mesenchyme of mandibular portion of arch one (1) and the circumpharyngeal region (arrow). **C-F.** Lateral (C, D) and ventral (E, F) views of E9.5 *Gata3^+/z^* (E, G) and *Gata3^z/z^* (F, H) embryos. In both genotypes, β-gal activity is observed in the mesenchyme and overlying ectoderm of the ventral mandibular arch. In *Gata3^z/z^* embryos, β-gal activity is also observed surrounding the optic placode (op) and maxillary prominence. **G-L.** Lateral (G, I), ventral (H, J) and transverse sectional views (K, L) of E10.5 *Gata3^+/z^* (E, G) and *Gata3^z/z^* (F, H) embryos. Sections are counterstained with eosin. In *Gata3^z/+^* embryos, prominent β-gal activity is confined to the ventral mandibular arch mesenchyme (red-dashed line in H, K; the arch is outlined in yellow in H) and overlying ectoderm (K). Staining is also strong in the medial (mnp) and lateral (lnp) nasal prominences at the lambdoidal (*λ*) junction. In E10.5 *Gata3^z/z^* embryos, staining is similar to that of *Gata3^+/z^* embryos (I, J, L), though strong staining extends dorsally from the distal staining domain in the mandibular arch (outlined by yellow-dashed line) along the rostral half of the arch (red-dashed line) into the maxillomandibular junction, where it meets the staining in the lnp. frontonasal prominence; h, heart; lb, limb bud; op, optic placode; ot, otic placode; otr, outflow tract; 2, second pharyngeal arch.

Specifically, in pharyngeal arch 1, staining was observed in the ventral half of the mandibular arch in both *Gata3^+/z^* (Fig. 8C, D) and *Gata3^z/z^* (Fig. 8E, F) embryos. Staining was also apparent in the otic placode and outflow tract of both genotypes (Fig. 8C-F).

At E10.5, *β*-gal staining in *Gata3^+/z^* embryos was still confined to the ventral mandibular arch (red dashed line in Fig. 8H-K), with staining present in the ectoderm and underlying mesenchyme of the arch (Fig. 8F). *β*-gal staining was also present in the posterior medial and lateral nasal prominences where they fuse (the lambdoidal junction, *λ*) (Fig. 8E). In E10.5 *Gata3^z/z^* embryos, however, *β*-gal activity in the ventral mandibular arch extended along the rostral half of the mandibular arch (red dashed line in Fig. 8J, L) from the ventral domain into the maxillomandibular junction (Fig, I, J, L), where it met the staining in the lateral nasal prominence next to the lambdoidal junction. These findings indicate that, like the findings from ISH, loss of GATA3 allows aberrant upregulation of a ventral GRN that includes *Gata3* itself between the mandibular arch ventral cap and the hinge. It thus appears that a key function of GATA3 is to maintain separation of patterning cues between the maxilla and mandibular portions of arch 1, ensuring separation of the upper and lower jaws.

## Discussion

We demonstrate here that GATA3 is crucial for craniofacial development, with loss both reducing the total NCC number in the early mandibular arch and disrupting the balance between BMP and FGF8 GRNs Together, our results illustrate that GATA3 is essential for separating the morphogenetic programs that define the upper and lower jaws, acting as a factor to ensure the maxillomandibular (hinge) region defined in the hinge and caps model develops without intrusion of cap signals.

### Early GATA3 function

Hypoplasia of numerous craniofacial structures in E18.5 *Gata3^z/z^* embryos is preceded by hypoplasia of the mandibular arch at E9.5. While cell proliferation and cell death are unchanged in E9.5 post-migratory NCCs, the total number of cells within the mandibular arch are significantly reduced. *Msx1^-/-^*/*Msx2*^-/-^ embryos also exhibit craniofacial bone hypoplasia that is preceded by arch hypoplasia at E9.5 (Ishii et al., 2005). However, while the expression of *Tfap2* is delayed in *Msx1^-/-^*/*Msx2*^-/-^ embryos, expression of *Tfap2* in *Gata3^+/+^* and *Gata3^z/z^* embryos between E8.5 and E9.5 appears temporally similar, though staining always appears weaker in *Gata3^z/z^* mutant embryos at this stage. GATA2/3 are key modulators of a transcriptional circuit that positions the neural plate border, and act directly downstream mediators of early BMP signaling [(Linker et al., 2009); reviewed in (Simões-Costa and Bronner, 2015)],. It is thus possible that loss of GATA3 shifts the boundary of the neural plate border, thus allowing an expansion of the neural plate or placodal domain at the expense of NCC, thus reducing the number of NCCs available to populate the pharyngeal arches. Further experiments are required to directly test this possibility.

### Loss of GATA3 disrupts GRNs around the maxillomandibular junction

One of the key signals establishing the maxillomandibular junction is FGF8. In chick embryos, Fgf8 in the early ectoderm defines the maxillomandibular junction, with *Bmp4* expression in the ventral ectoderm preventing a more ventral expansion *Fgf8* expression (Shigetani et al., 2000).

Targeted inactivation of *Fgf8* using *Nestin-Cre* in mouse embryos leads to a dramatic reduction in the size of the mandibular arch and NCC-derived mesenchyme cell death, suggesting that the patterning program induced by ectodermal FGF8 signaling in the pharyngeal arch is either directly or indirectly required for mesenchyme cell survival (Trumpp et al., 1999). However, in *Fgf8;Nes-Cre* mutant embryos, early *Bmp4* expression does not spread dorsally, indicating that in *Gata3^z/z^* embryos (Trumpp et al., 1999), downregulation of *Fgf8* expression in the maxillomandibular junction is not responsible for the upregulated *Bmp4* expression. In fact, that *Pitx2* expression is not dramatically changed in *Gata3^z/z^* embryos suggests the opposite: changes in the *Fgf8* GRN occur due to an expansion of BMP signaling. BMP4 bead implantation in mandibular arch explants represses *Fgf8* expression, indicating that BMP4 may more directly repress *Fgf8* transcription (Stottmann et al., 2001). Similarly, targeted expression of *Bmp4* in mouse NCCs also leads to downregulation of *Fgf8* and an upregulation in the expression of BMP-responsive genes, including *Hand2* and *Msx1*, resulting in bony syngnathia (He et al., 2014). Further, BMP4 overexpression was shown to induce a BIG (<u>B</u>MP induced <u>g</u>ene) profile that includes *Hand2*, *Gata3*, *Msx1* and *Hand1* (Bonilla-Claudio et al., 2012). In contrast, loss of *Bmp4* within the arch ectoderm led to a reduction in the expression of these BIG genes (except for *Hand1*; discussed below). Thus, changes in the BIG profile appear to be a major driver of the gene expression changes in *Gata3^z/z^* embryos.

If loss of GATA3 results in the inappropriate upregulation of a BMP GRN within the maxillomandibular junction, what then is responsible for syngnathia? Several proteins have been demonstrated to be involved in jaw separation and whose disruption also leads to syngnathia, one of which is the gene encoding FOXC1, a forkhead box winged helix transcription factor that is expressed at E8.5 within the ectoderm and mesenchyme of the first arch (Inman et al., 2013). Disruption of *Foxc1* leads to syngnathia, with variations in phenotype involving a genetic interaction between *Fgf8* and *Foxc1* (Inman et al., 2013). This is similar to the syngnathia observed following conditional expression of *Bmp4* in NCCs described above (He et al., 2014). These results suggest that while aberrant upregulation of a *Bmp4* GRN in the maxillomandibular junction leads to syngnathia, it is the subsequent downregulation of a *Fgf8* GRN that derails boundary separation between the mandibular and maxillary prominences and hence syngnathia.

### Confinement of gene expression in the mandibular ventral cap

One obvious requirement of the Hinge and Caps model is that ventral cap gene expression must be confined to the ventral portions of the mandibular arch to ensure normal patterning of the dorsal arch. One example of this is *Hand1*, whose tightly restricted ventral cap expression requires the direct overlap of BMP signaling and HAND2 transcriptional activity (Vincentz et al., 2016). However, while *Hand2* expression in *Gata3^z/z^* embryos spreads in a dorsorostral direction towards the maxillomandibular junction (Ruest et al., 2004), *Hand1* expression does not. This illustrates that while BMP and HAND2 are both required for *Hand1* expression (Barron et al., 2011; Vincentz et al., 2016), repressive mechanisms also exist that can override these inductive mechanisms. The *Hand1* pharyngeal arch enhancer [Hand1^PA/OFT^ enhancer; (Vincentz et al., 2016)] contains several validated DLX binding elements, and introduction of DLX5 in reporter assays disrupts BMP/HAND2 synergy *in vitro*, thus preventing expression of *Hand1^PA/OFT^* enhancer-driven transgenes (Vincentz et al., 2016). Given that *Dlx5* expression in *Gata3* mutant embryos expands in a pattern similar to *Hand2* (Ruest et al., 2004), the DLX repressive activity likely prevents dorsal *Hand1* expansion. A targeted deletion approach utilizing CRISPR to mutagenize DLX DNA binding consensus residues is required to address this question.

### *Gata3* loss-of-function as a model for Hemifacial microsomia and development of lateral symmetry

We have shown here that loss of *Gata3* leads to a disruption in lateral symmetry; while both sides of the facial skeleton, including the mandible, maxilla and middle ear structures are affected in *Gata3^z/z^* embryos, one side is always more severely affected than the other. These changes resemble those in human individuals with Hemifacial microsomia (HFM; MIM 164210; also known as Goldenhar syndrome or oculoauriculovertebral spectrum), which often includes unilateral defects in first and second pharyngeal arch-arch derived structures, including the mandible, temporomandibular joint, middle ear bone, ear pinna, maxilla, zygoma and muscles of mastication (Tiner and Quaroni, 1996; Barisic et al., 2014). The vast majority of HFM individuals do not have a genetic diagnosis. While *BAPX1* (Fischer et al., 2006), *MYT1* (Lopez et al., 2016; Berenguer et al., 2017), *TCOF1* and *SALL1* (Huang et al., 2010) have been implicated in HFM individuals, none have been validated (Thiel et al., 2005; Fischer et al., 2006). Further, mouse mutants for *Bapx1* (Tribioli and Lufkin, 1999; Akazawa et al., 2000; Tucker et al., 2004), *Myt1* (Wang et al., 2007) and *Tcof1* (Dixon et al., 2006) have not been reported to have unilateral craniofacial defects. One of the few mouse models reported as a model for HFM resulted from a transgenic insertional event on mouse chromosome 10 (Naora et al., 1994; Cousley et al., 2002). While this locus was named the Hemifacial microsomia-associated (*Hfm*) locus, the underlying genetic lesion has not been described. *Goosecoid* (*Gsc*) is located within the insertional region, though mutations in *GSC* were not found in two groups of HFM individuals (Kelberman et al., 2001). In addition, *Gsc^-/-^* mouse embryos do not develop a *Hfm* phenotype (Rivera-Perez et al., 1995; Yamada et al., 1995; Rivera-Perez et al., 1999). Our results here provide a new gene to explore in HFM individuals. Indeed, Genome Wide Association Studies have identified the locus containing *GATA3* as a susceptibility locus for craniofacial microsomia (p=6.58 × 10^−9^) (Zhang et al., 2016).

## Acknowledgements

The authors would like to thank Katherine Kuhn and Drs. Andre Tavares and Gwinn Vonnahme for technical assistance and scientific input and Dr. Katherine Fantauzzo for critical reading, suggestions and encouragement.

## REFERENCES

1. Abe, M., Ruest, L.-B. and Clouthier, D. E. (2007). Fate of cranial neural crest cells during craniofacial development in endothelin-A receptor deficient mice. Int. J. Dev. Biol. 51, 97–105.

2. Akazawa, H., Komuro, I., Sugitani, Y., Yazaki, Y., Nagai, R. and Noda, T. (2000). Targeted disruption of the homeobox transcription factor Bapx1 results in lethal skeletal dysplasia with asplenia and gastroduodenal malformation. Genes Cell 5, 499–513.

3. Barisic, I., Odak, L., Loane, M., Garne, E., Wellesley, D., Calzolari, E., Dolk, H., Addor, M. C., Arriola, L., Bergman, J. et al. (2014). Prevalence, prenatal diagnosis and clinical features of oculo-auriculo-vertebral spectrum: a registry-based study in Europe. Eur J Hum Genet 22, 1026–1033.

4. Barron, F., Woods, C., Kuhn, K., Bishop, J., Howard, M. J. and Clouthier, D. E. (2011). Downregulation of *Dlx5* and *Dlx6* expression by Hand2 is essential for initiation of tongue morphogenesis. Development 138, 2249–2259.

5. Berenguer, M., Tingaud-Sequeira, A., Colovati, M., Melaragno, M. I., Bragagnolo, S., Perez, A. B. A., Arveiler, B., Lacombe, D. and Rooryck, C. (2017). A novel de novo mutation in MYT1, the unique OAVS gene identified so far. Eur J Hum Genet 25, 1083–1086.

6. Bonilla-Claudio, M., Wang, J., Bai, Y., Klysik, E., Selever, J. and Martin, J. F. (2012). Bmp signaling regulates a dose-dependent transcriptional program to control facial skeletal development. Development 139, 709–719.

7. Brewer, S., Feng, W., Huang, J., Sullivan, S. and Williams, T. (2004). Wnt1-Cre-mediated deletion of AP-2alpha causes multiple neural crest-related defects. Dev Biol 267, 135–152.

8. Charité, J., McFadden, D. G., Merlo, G. R., Levi, G., Clouthier, D. E., Yanagisawa, M., Richardson, J. A. and Olson, E. N. (2001). Role of Dlx6 in regulation of an endothelin-1-dependent, *dHAND* branchial arch enhancer. Genes Dev. 15, 3039–3049.

9. Clouthier, D. E., Garcia, E. and Schilling, T. F. (2010). Regulation of facial morphogenesis by endothelin signaling: insights from mouse and fish. *Am. J. Med. Genet.*, Part A. 152A, 2962-2973.

10. Clouthier, D. E., Williams, S. C., Yanagisawa, H., Wieduwilt, M., Richardson, J. A. and Yanagisawa, M. (2000). Signaling pathways crucial for craniofacial development revealed by endothelin-A receptor-deficient mice. Dev. Biol. 217, 10–24.

11. Clouthier, D. E., Passos-Bueno, M. R., Tavares, A. L., Lyonnet, S., Amiel, J. and Gordon, C. T. (2013). Understanding the basis of auriculocondylar syndrome: Insights from human, mouse and zebrafish genetic studies. Am J Med Genet C Semin Med Genet 163c, 306-317.

12. Clouthier, D. E., Hosoda, K., Richardson, J. A., Williams, S. C., Yanagisawa, H., Kuwaki, T., Kumada, M., Hammer, R. E. and Yanagisawa, M. (1998). Cranial and cardiac neural crest defects in endothelin-A receptor-deficient mice. Development 125, 813–824.

13. Cousley, R., Naora, H., Yokoyama, M., Kimura, M. and Otani, H. (2002). Validity of the Hfm transgenic mouse as a model for hemifacial microsomia. Cleft Palate Craniofac J 39, 81–92.

14. Dee, C. T., Szymoniuk, C. R., Mills, P. E. and Takahashi, T. (2013). Defective neural crest migration revealed by a Zebrafish model of Alx1-related frontonasal dysplasia. Hum Mol Genet 22, 239–251.

15. Depew, M. J. and Compagnucci, C. (2008). Tweaking the hinge and caps: testing a model of the organization of jaws. J. Exp. Zool. (Mol. Dev. Evol.) 310, 315–335.

16. Depew, M. J., Lufkin, T. and Rubenstein, J. L. (2002). Specification of jaw subdivisions by *Dlx* genes. Science 298, 381–385.

17. Depew, M. J., Simpson, C. A., Morasso, M. and Rubenstein, J. L. R. (2005). Reassessing the *Dlx* code: the genetic regulation of branchial arch skeletal pattern and development. J. Anat. 207, 501–561.

18. Dixon, J., Jones, N. C., Sandell, L. L., Jayasinghe, S. M., Crane, J., Rey, J. P., Dixon, M. J. and Trainor, P. (2006). *Tcof1*/Treacle is required for neural crest cell formation and proliferation deficiencies that cause craniofacial abnormalities. Proc. Natl. Acad. Sci. U S A 103, 13403–13408.

19. Ferguson, C. A., Tucker, A. S. and Sharpe, P. T. (2000). Temporospatial cell interactions regulating mandibular and maxillary arch patterning. Development 127, 403–412.

20. Fischer, S., Lüdecke, H. J., Wieczorek, D., Böhringer, S., Gillessen-Kaesbach, G. and Horsthemke, B. (2006). Histone acetylation dependent allelic expression imbalance of BAPX1 in patients with the oculo-auriculo-vertebral spectrum. Hum Mol Genet 15, 581–587.

21. Grote, D., Boualia, S. K., Souabni, A., Merkel, C., Chi, X., Costantini, F., Carroll, T. and Bouchard, M. (2008). Gata3 acts downstream of beta-catenin signaling to prevent ectopic metanephric kidney induction. PLoS Genet 4, e1000316.

22. He, F., Hu, X., Xiong, W., Li, L., Lin, L., Shen, B., Yang, L., Gu, S., Zhang, Y. and Chen, Y. (2014). Directed Bmp4 expression in neural crest cells generates a genetic model for the rare human bony syngnathia birth defect. Dev Biol 391, 170–181.

23. Hendershot, T. J., Liu, H., Clouthier, D. E., Shepherd, I. T., Coppola, E., Studer, M., Firulli, A. B., Pittman, D. L. and Howard, M. J. (2008). Conditional deletion of *Hand2* reveals critical functions in neurogenesis and cell type-specific gene expression for development of neural crest-derived noradrenergic sympathetic ganglion neurons. Dev. Biol. 319, 179–191.

24. Ho, I. C. and Pai, S. Y. (2007). GATA-3 - not just for Th2 cells anymore. Cell Mol Immunol 4, 15–29.

25. Huang, X. S., Li, X., Tan, C., Xiao, L., Jiang, H. O., Zhang, S. F., Wang, D. M. and Zhang, J. X. (2010). Genome-wide scanning reveals complex etiology of oculo-auriculo-vertebral spectrum. Tohoku J Exp Med 222, 311–318.

26. Inman, K. E., Purcell, P., Kume, T. and Trainor, P. A. (2013). Interaction between Foxc1 and Fgf8 during mammalian jaw patterning and in the pathogenesis of syngnathia. PLoS Genet 9, e1003949.

27. Ishii, M., Han, J., Yen, H.-Y., Sucov, H. M., Chai, Y. and Maxson, J., R.E. (2005). Combined deficiencies of *Msx1* and *Msx2* cause impaired patterning and survival of the cranial neural crest. Development 132, 4937–4950.

28. Kelberman, D., Tyson, J., Chandler, D. C., McInerney, A. M., Slee, J., Albert, D., Aymat, A., Botma, M., Calvert, M., Goldblatt, J. et al. (2001). Hemifacial microsomia: progress in understanding the genetic basis of a complex malformation syndrome. Hum Genet 109, 638–645.

29. Lakshmanan, G., Lieuw, K. H., Lim, K.-C., Gu, Y., Grosveld, F., Engel, J. D. and Karis, A. (1999). Localization of distant urogenital system-, central nervous system-, and endocardium-specific transcriptional regulatory elements in the GATA-3 locus. Mol Cell Biol 19, 1558–1568.

30. Lanctôt, C., Lamolet, B. and Drouin, J. (1997). The bicoid-related homeoprotein Ptx1 defines the most anterior domain of the embryo and differentiates posterior from anterior lateral mesoderm. Development 124, 2807–2817.

31. Lim, K.-C., Lakshmanan, G., Crawford, S. E., Gu, Y., Grosveld, F. and Engel, J. D. (2000). *Gata3* loss leads to embryonic lethality due to noradrenaline deficiency of the sympathetic nervous system. Nat Gen 25, 209–212.

32. Linker, C., De Almeida, I., Papanayotou, C., Stower, M., Sabado, V., Ghorani, E., Streit, A., Mayor, R. and Stern, C. D. (2009). Cell communication with the neural plate is required for induction of neural markers by BMP inhibition: evidence for homeogenetic induction and implications for Xenopus animal cap and chick explant assays. Dev Biol 327, 478–486.

33. Liu, W., Selever, J., Lu, M.-F. and Martin, J. F. (2003a). Genetic dissection of *Pitx2* in craniofacial development uncovers new functions in branchial arch morphogenesis, late aspects of tooth morphogenesis and cell migration. Development 130, 6375–6385.

34. Liu, W., Selever, J., Lu, M. F. and Martin, J. F. (2003b). Genetic dissection of Pitx2 in craniofacial development uncovers new functions in branchial arch morphogenesis, late aspects of tooth morphogenesis and cell migration. Development 130, 6375–6385.

35. Liu, W., Selever, J., Murali, D., Sun, X., Brugger, S. M., Ma, L., Schwartz, R. J., Maxson, R., Furuta, Y. and Martin, J. F. (2005). Threshold-specific requirements for Bmp4 in mandibular development. Dev Biol 283, 282–293.

36. Lopez, E., Berenguer, M., Tingaud-Sequeira, A., Marlin, S., Toutain, A., Denoyelle, F., Picard, A., Charron, S., Mathieu, G., de Belvalet, H. et al. (2016). Mutations in MYT1, encoding the myelin transcription factor 1, are a rare cause of OAVS. J Med Genet 53, 752–760.

37. Maeda, A., Moriguchi, T., Hamada, M., Kusakabe, M., Fujioka, Y., Nakano, T., Yoh, K., Lim, K. C., Engel, J. D. and Takahashi, S. (2009). Transcription factor GATA-3 is essential for lens development. Dev Dyn 238, 2280–2291.

38. Medeiros, D. M. and Crump, J. G. (2012). New perspectives on pharyngeal dorsoventral patterning in development and evolution of the vertebrate jaw. Dev Biol 371, 121–135.

39. Naora, H., Kimura, M., Otani, H., Yokoyama, M., Koizumi, T., Katsuki, M. and Tanaka, O. (1994). Transgenic mouse model of hemifacial microsomia: cloning and characterization of insertional mutation region on chromosome 10. Genomics 23, 515–519.

40. Neben, C. L. and Merrill, A. E. (2015). Signaling Pathways in Craniofacial Development: Insights from Rare Skeletal Disorders. Curr Top Dev Biol 115, 493–542.

41. Pandolfi, P. P., Roth, M. E., Karis, A., Leonard, M. W., Dzierzak, E., Grosveld, F., Engel, J. D. and Lindenbaum, M. H. (1995). Targeted deletion of the *GATA3* gene causes severe abnormalities in the nervous system and in fetal liver haematopoiesis. Nat Gen 11, 40–44.

42. Raid, R., Krinka, D., Bakhoff, L., Abdelwahid, E., Jokinen, E., Kärner, M., Malva, M., Meier, R., Pelliniemi, L. J., Ploom, M. et al. (2009). Lack of Gata3 results in conotruncal heart anomalies in mouse. Mech Dev 126, 80–89.

43. Rivera-Perez, J. A., Wakamiya, M. and Behringer, R. R. (1999). *Goosecoid* acts cell autonomously in mesenchyme-derived tissues during craniofacial development. Development 126, 3811–3821.

44. Rivera-Perez, J. A., Mallo, M., Gendron-Maguire, M., Gridley, T. and Behringer, R. R. (1995). *goosecoid* is not an essential component of the mouse gastrula organizer but is required for craniofacial and rib development. Development 121, 3005–3012.

45. Ruest, L. B., Xiang, X., Lim, K. C., Levi, G. and Clouthier, D. E. (2004). Endothelin-A receptor-dependent and -independent signaling pathways in establishing mandibular identity. Development 131, 4413–4423.

46. Sato, T., Kawamura, Y., Asai, R., Amano, T., Uchijima, Y., Dettlaff-Swiercz, D. A., Offermanns, S., Kurihara, Y. and Kurihara, H. (2008). Recombinase-mediated cassette exchange reveals the selective use of Gq/G11-dependent and -independent endothelin 1-endothelin type A receptor signaling in pharyngeal arch development. Development 135, 755–765.

47. Shigetani, Y., Nobusada, Y. and Kuratani, S. (2000). Ectodermally derived FGF8 defines the maxillomandibular region in the early chick embryo: epithelial-mesenchymal interactions in the specification of the craniofacial ectomesenchyme. Dev Biol 228, 73–85.

48. Simões-Costa, M. and Bronner, M. E. (2015). Establishing neural crest identity: a gene regulatory recipe. Development 142, 242–257.

49. Stottmann, R. W., Anderson, R. M. and Klingensmith, J. (2001). The BMP antagonists Chordin and Noggin have essential but redundant roles in mouse mandibular outgrowth. Dev Biol 240, 457–473.

50. Tavares, A. L. and Clouthier, D. E. (2015). Cre recombinase-regulated Endothelin1 transgenic mouse lines: novel tools for analysis of embryonic and adult disorders. Dev Biol 400, 191–201.

51. Tavares, A. L. P., Cox, T. C., Maxson, R. M., Ford, H. L. and Clouthier, D. E. (2017). Negative regulation of Endothelin signaling by SIX1 is required for proper maxillary development. Development 144, 2021–2031.

52. Thiel, C. T., Rosanowski, F., Kohlhase, J., Reis, A. and Rauch, A. (2005). Exclusion of TCOF1 mutations in a case of bilateral Goldenhar syndrome and one familial case of microtia with meatal atresia. Clin Dysmorphol 14, 67–71.

53. Tiner, B. D. and Quaroni, A. L. (1996). Facial asymmetries in hemifacial microsomia, Goldenhar syndrome, and Treacher Collins syndrome. Atlas Oral Maxillofac Surg Clin North Am 4, 37–52.

54. Tissier-Seta, J.-P., Mucchielli, M.-L., Mark, M., M.-G., Goridis, C. and Brunet, J.-F. (1995). *Barx1*, a new mouse homeodomain transcription factor expressed in cranio-facial ectomesenchyme and the stomach. Mech Dev 51, 3–15.

55. Tribioli, C. and Lufkin, T. (1999). The murine Bapx1 homeobox gene plays a critical role in embryonic development of the axial skeleton and spleen. Development 126, 5699–5711.

56. Trummp, A., Depew, M. J., Rubenstein, J. L. R., Bishop, J. M. and Martin, G. R. (1999). Cre-mediated gene inactivation demonstrates that FGF8 is required for cell survival and patterning of the first branchial arch. Genes Dev 13, 3136–3148.

57. Trumpp, A., Depew, M. J., Rubenstein, J. L., Bishop, J. M. and Martin, G. R. (1999). Cre-mediated gene inactivation demonstrates that FGF8 is required for cell survival and patterning of the first branchial arch. Genes Dev 13, 3136–3148.

58. Tucker, A. S., Watson, R. P., Lettice, L. A., Yamada, G. and Hill, R. E. (2004). Bapx1 regulates patterning in the middle ear: altered regulatory role in the transition from the proximal jaw during vertebrate evolution. Development 131, 1235–1245.

59. Tucker, S. A., Yamada, G., Grigoriou, M., Pachnis, V. and Sharpe, P. T. (1999). Fgf-8 determines rostral-caudal polarity in the first branchial arch. Development 126, 51–61.

60. Uz, E., Alanay, Y., Aktas, D., Vargel, I., Gucer, S., Tuncbilek, G., von Eggeling, F., Yilmaz, E., Deren, O., Posorski, N. et al. (2010). Disruption of ALX1 causes extreme microphthalmia and severe facial clefting: expanding the spectrum of autosomal-recessive ALX-related frontonasal dysplasia. Am J Hum Genet 86, 789–796.

61. Van Esch, H. and Devriendt, K. (2001). Transcription factor GATA3 and the human HDR syndrome. Cell Mol Life Sci 58, 1296–1300.

62. Vincentz, J. W., Casasnovas, J. J., Barnes, R. M., Que, J., Clouthier, D. E., Wang, J. and Firulli, A. B. (2016). Exclusion of *Dlx5/6* expression from the distal-most mandibular arches enables BMP-mediated specification of the distal cap. Proc Natl Acad Sci U S A 113, 7563–7568.

63. Wang, S., Zhang, J., Zhao, A., Hipkens, S., Magnuson, M. A. and Gu, G. (2007). Loss of Myt1 function partially compromises endocrine islet cell differentiation and pancreatic physiological function in the mouse. Mech Dev 124, 898–910.

64. Yamada, G., Mansouri, A., Torres, M., Stuart, E. T., Blum, M., Schultz, M., De Robertis, E. M. and Gruss, P. (1995). Targeted mutation of the murine *goosecoid* gene results in craniofacial defects and neonatal death. Development 121, 2917–2922.

65. Zhang, Y. B., Hu, J., Zhang, J., Zhou, X., Li, X., Gu, C., Liu, T., Xie, Y., Liu, J., Gu, M. et al. (2016). Genome-wide association study identifies multiple susceptibility loci for craniofacial microsomia. Nat Commun 7, 10605.

